# Acitretin mitigates uroporphyrin-induced bone defects in congenital erythropoietic porphyria models

**DOI:** 10.1101/2021.03.30.437658

**Authors:** Juliana Bragazzi Cunha, Jared S Elenbaas, Dhiman Maitra, Ning Kuo, Rodrigo Azuero-Dajud, Allison C Ferguson, Megan S Griffin, Stephen I Lentz, Jordan A Shavit, M Bishr Omary

## Abstract

Congenital erythropoietic porphyria (CEP) is a rare genetic disorder leading to accumulation of uro/coproporphyrin-I in tissues due to inhibition of uroporphyrinogen-III synthase. Clinical manifestations of CEP include bone fragility, severe photosensitivity and photomutilation. Currently there is no specific treatment for CEP, except bone marrow transplantation, and there is an unmet need for treating this orphan disease. Fluorescent porphyrins cause protein aggregation, which led us to hypothesize that uroporphyrin-I accumulation leads to protein aggregation and CEP-related bone phenotype. We developed a zebrafish model that phenocopies features of CEP. As in human patients, uroporphyrin-I accumulated in the bones of zebrafish, leading to impaired bone development. Furthermore, in an osteoblast-like cell line, uroporphyrin-I decreased mineralization, aggregated bone matrix proteins, activated endoplasmic reticulum stress and disrupted autophagy. Using high-throughput drug screening, we identified acitretin, a second-generation retinoid, and showed that it reduced uroporphyrin-I accumulation and its deleterious effects on bones. Our findings provide a new CEP experimental model and a potential repurposed therapeutic.

## INTRODUCTION

Porphyrias are a group of inherited disorders due to defects in the heme biosynthetic pathway^1,2^. One such example is congenital erythropoietic porphyria (CEP), most commonly caused by loss of function mutation in uroporphyrinogen III synthase (UROS; Fig.S1), the enzyme that catalyzes the third step of the heme biosynthetic pathway^1,3^. CEP is rare, with ~250 cases reported to date^4,5^. It is autosomal recessive, and associated with reduced UROS activity (5% of normal) and consequent accumulation of uro/coproporphyrin-I (uro/copro-I) in bone marrow, erythrocytes, plasma, and increased uro/copro-I excretion in urine and stool^1,3,5,6^. CEP is characterized by severe photosensitivity, with skin fragility and blistering of sun-exposed areas^1,5,6^. Scaring due to secondary skin infections and bone resorption contribute to disfigurement of light-exposed areas^3^. Other clinical manifestations of this multisystem disease include chronic ulcerative keratitis, hemolysis, which may require repeated blood transfusions in severe cases, nonimmune hydrops fetalis, red urine since birth, erythrodontia and osteodystrophy^1,3,6,7^. Currently, there is no specific pharmacological treatment for CEP, with interventions being life-style-related (e.g. avoidance of sun) or complex procedures, including bone marrow transplantation^3,8,9^.

Fluorescent porphyrin accumulation in porphyria causes organelle specific protein oxidation and aggregation through mechanisms that involve type-II photosensitive reactions and secondary oxidative stress^10–17^. We posit that porphyrin-mediated protein aggregation in CEP plays a major mechanistic role in tissue damage that involves accumulation of fluorescent uro/copro-I.

We demonstrated that uro-I injection of zebrafish larvae mimics features of CEP, including uro-I accumulation in bones and bone deformation, as judged by decreased vertebra and operculum volume. Uro-I treatment of an osteoblastic human osteosarcoma cell line, Saos-2, caused significant decrease in mineral matrix synthesis and proteotoxicity. Using high-throughput drug screening, we identified acitretin, a 2^nd^ generation retinoid, as an effective drug that mitigates some of the harmful effects of uro-I in zebrafish and Saos-2 cells.

## RESULTS

### An inducible zebrafish model mimics bone defects of human CEP

Uro-I injected zebrafish larvae showed porphyrin fluorescence in bone tissue (Fig.1A). To confirm that uro-I binds specifically to bone, larvae were co-injected with calcein (bone-specific dye) and imaged. Calcein and uro-I fluorescence co-localized (Fig.1B), confirming uro-I bound to bone. Additionally, uro-I-injected larvae exhibited severe photosensitivity and had to be shielded from light to prevent their death (not shown). Next, we assessed uro-I-mediated bone defect by measuring the volume of the operculum and 4^th^ vertebra. Notably, uro-I injection significantly decreased operculum and 4^th^ vertebra volume (Fig.1C).

**Figure 1.**
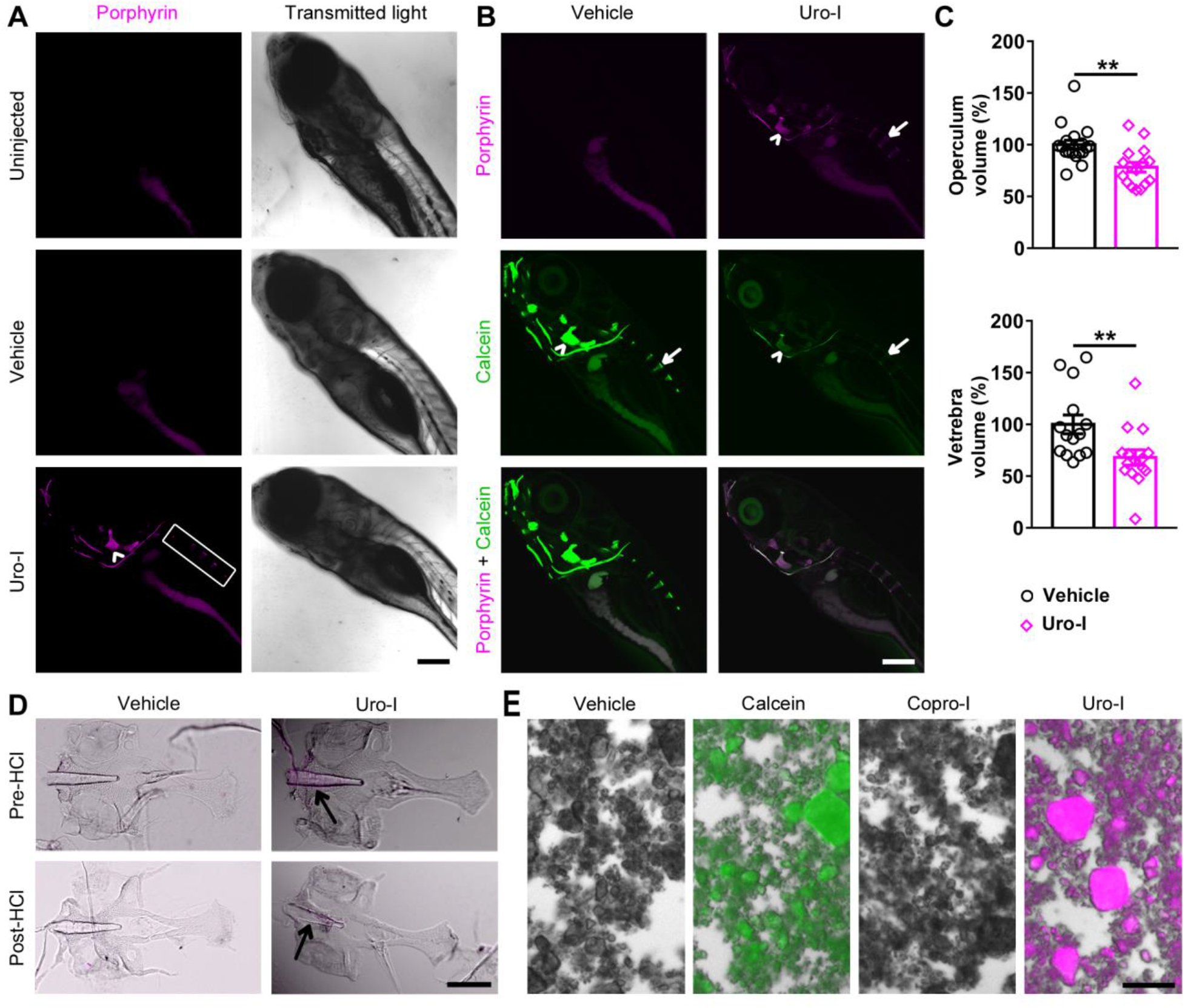
Zebrafish model of CEP develops bone phenotype resembling human disease. **(A)** 6dpf zebrafish larvae were injected with uro-I or vehicle and imaged by confocal microscopy at 7dpf. Porphyrin was detected only in the bones of uro-I-injected group. Arrowhead-operculum; box-vertebrae. **(B)** Larvae were treated as in (A) and injected with calcein prior to imaging. Arrowhead-operculum; arrow-4^th^ vertebra. **(C)** Quantification of bone volume in larvae from (B); bone volume was normalized to vehicle-injected larvae set to 100%. Symbols represent individual larvae (14-18/group) from 4-5 independent experiments. **(D)** Larvae were treated as in (A). At 7dpf bones were harvested and imaged by epifluorescence microscopy pre and post HCl bone demineralization, arrow-notochord. **(E)** Hydroxyapatite was incubated with calcein/uro-I/copro-I and imaged by epifluorescence microscopy. Scale bars: 200μm (A-D); 50μm (E). \*\**p*<0.01

Bone matrix is composed of protein/organic (including collagen/fibronectin/osteonectin) and inorganic components (minerals, mostly hydroxyapatite)^18,19^. We tested whether uro-I binds to the protein/organic or inorganic parts of bone matrix by demineralizing bones of uro-I-injected larvae. Demineralization caused loss of uro-I fluorescence, indicating that uro-I is extractable from the mineral matrix (Fig.1D). We validated this finding *in vitro* using hydroxyapatite crystals. Uro-I, but not copro-I, bound to hydroxyapatite, with calcein binding used as a positive control (Fig.1E). Therefore, uro-I binds to the inorganic bone matrix and its administration to zebrafish phenocopies three major features of CEP: osteal accumulation, bone defects, and severe photosensitivity.

### Acitretin mitigates uro-I effects in bones of zebrafish larvae

To identify potential drugs to treat CEP, we used our zebrafish CEP model and performed high-throughput screening of 1,280 small molecules by co-administering drug and uro-I injection (Fig.S2). Acitretin, a second-generation retinoid commonly used to treat psoriasis, decreased uro-I accumulation in bones (not shown). We validated the screening results and further characterized acitretin as a potential treatment for CEP by testing whether it had a prophylactic effect. Uro-I injected larvae were immediately transferred to either acitretin- or vehicle-containing medium. After 24h, acitretin-treated larvae had significantly reduced porphyrin fluorescence in their bones (operculum and vertebrae, Fig.2A,B) and increased uro-I excretion into the medium (Fig.2C), but no effect on operculum volume (Fig.2D). To assess the therapeutic potential of acitretin, larvae were injected with uro-I then transferred after 24h to medium containing either acitretin or vehicle and incubated for further 24h then imaged. Acitretin did not decrease bone porphyrin fluorescence (Fig.2E,F), but it increased uro-I excretion to the medium and operculum volume (Fig.2G,H).

**Figure 2.**
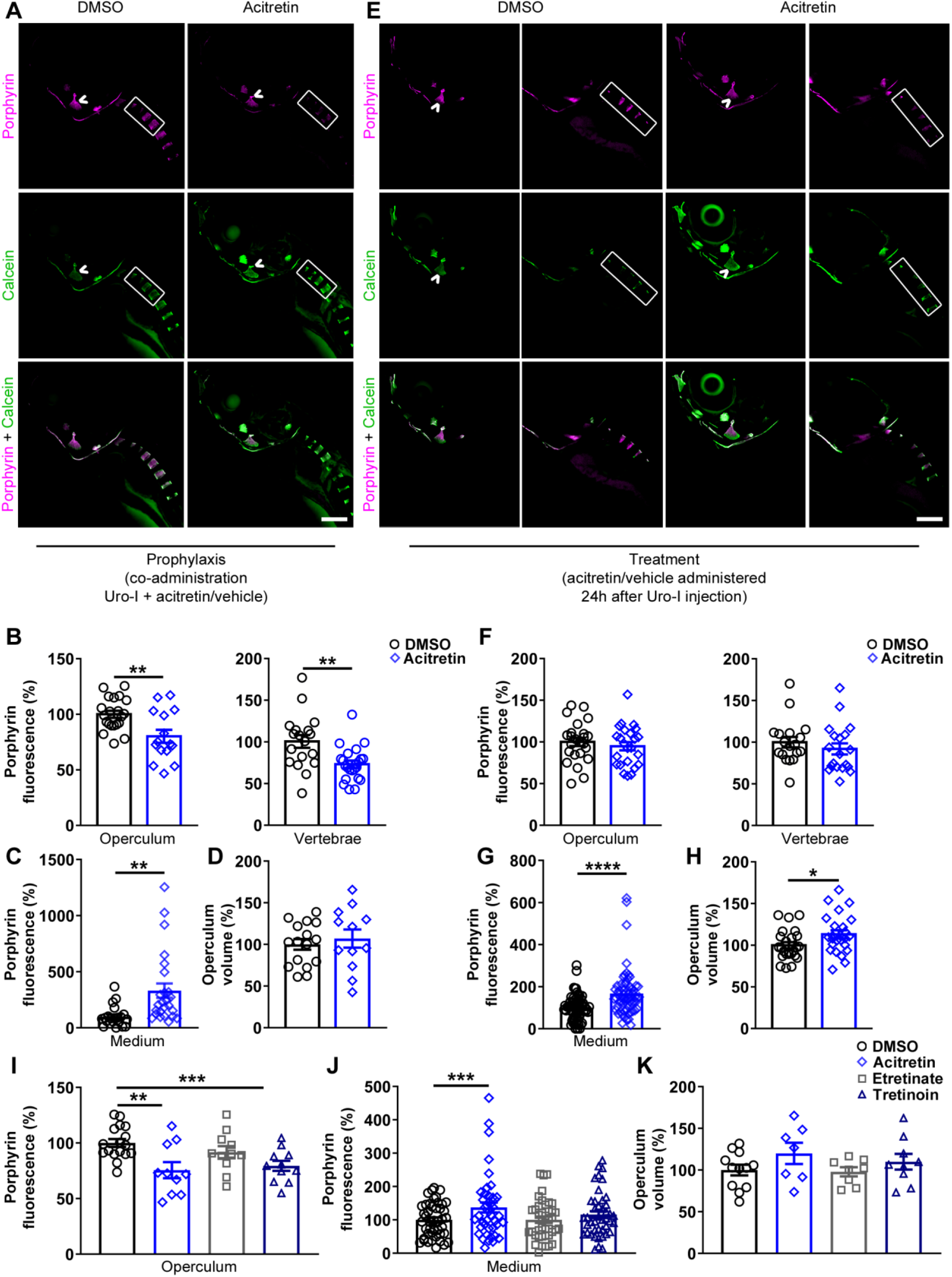
Acitretin mitigates CEP bone phenotype in zebrafish. **(A)** 6dpf larvae were injected with uro-I and transferred to medium containing acitretin or DMSO. At 7dpf larvae were injected with calcein and imaged by confocal microscopy. Quantification of porphyrin fluorescence **(B)**, porphyrin excretion **(C)** and operculum volume **(D)** from experiment in (A). Symbols represent individual larvae (12-25/group) from 3-4 independent experiments. **(E)** 6dpf larvae were injected with uro-I. At 7dpf they were transferred to medium containing acitretin or DMSO. At 8dpf larvae were injected with calcein and imaged by confocal microscopy. Quantification of porphyrin fluorescence **(F)**, porphyrin excretion **(G)** and operculum volume **(H)** from experiment in (E). Arrowhead-operculum; box-vertebrae (A,E). Symbols represent individual larvae (18-64/group) from 3-4 independent experiments. **(I,J,K)** Larvae were treated as in (A) with the indicated retinoid or DMSO and porphyrin fluorescence (I), porphyrin excretion (J) and operculum volume (K) were assayed. Bone volume was normalized to DMSO-treated larvae set to 100%, (D,H,K). Porphyrin excretion was normalized to DMSO-treated larvae set to 100%, (C,G,J). Symbols represent individual larvae (7-44/group) from 2-4 independent experiments. Scale bars: 200μm. \**p*<0.05, \*\**p*<0.01, \*\*\**p*<0.001, \*\*\*\**p*<0.0001

Since acitretin is a retinoid, we tested whether the protective effects are specific to acitretin or shared by other retinoids. Etretinate (second-generation retinoid and precursor of acitretin), tretinoin (all trans-retinoic acid) or acitretin were co-administered to zebrafish with uro-I. In addition to acitretin, tretinoin significantly reduced bone porphyrin (Fig.2I). Unlike acitretin, etretinate and tretinoin did not increase uro-I excretion into the medium (Fig.2J), and retinoids did not prevent loss in bone volume (Fig.2K). Thus, both retinoids, acitretin and tretinoin, prevent uro-I accumulation in zebrafish larvae. Hence, using our novel zebrafish CEP model, we demonstrated that acitretin attenuated uro-I-mediated bone damage by modulating the dynamics of uro-I bone binding and excretion.

### Uro-I impairs osteoblastic mineralization by aggregating matrix proteins, promoting ER stress and inhibiting autophagy

To elucidate the molecular mechanism of uro-I-mediated bone damage, we used Saos-2 cells, a human osteosarcoma cell line with osteoblastic features^20^. Mineralization was stimulated by treating cells with a mineralization activation cocktail (MAC) and assayed by alizarin red S (ARS) staining. As expected, Saos-2 cells manifested a mineralization phenotype when cultured for 3 days in MAC-supplemented medium, while in uro-I+MAC supplemented medium, mineralization decreased significantly (Fig.3A,B). Uro-I also caused marked photosensitivity, leading to cell death when cells were not shielded from light (not shown).

**Figure 3.**
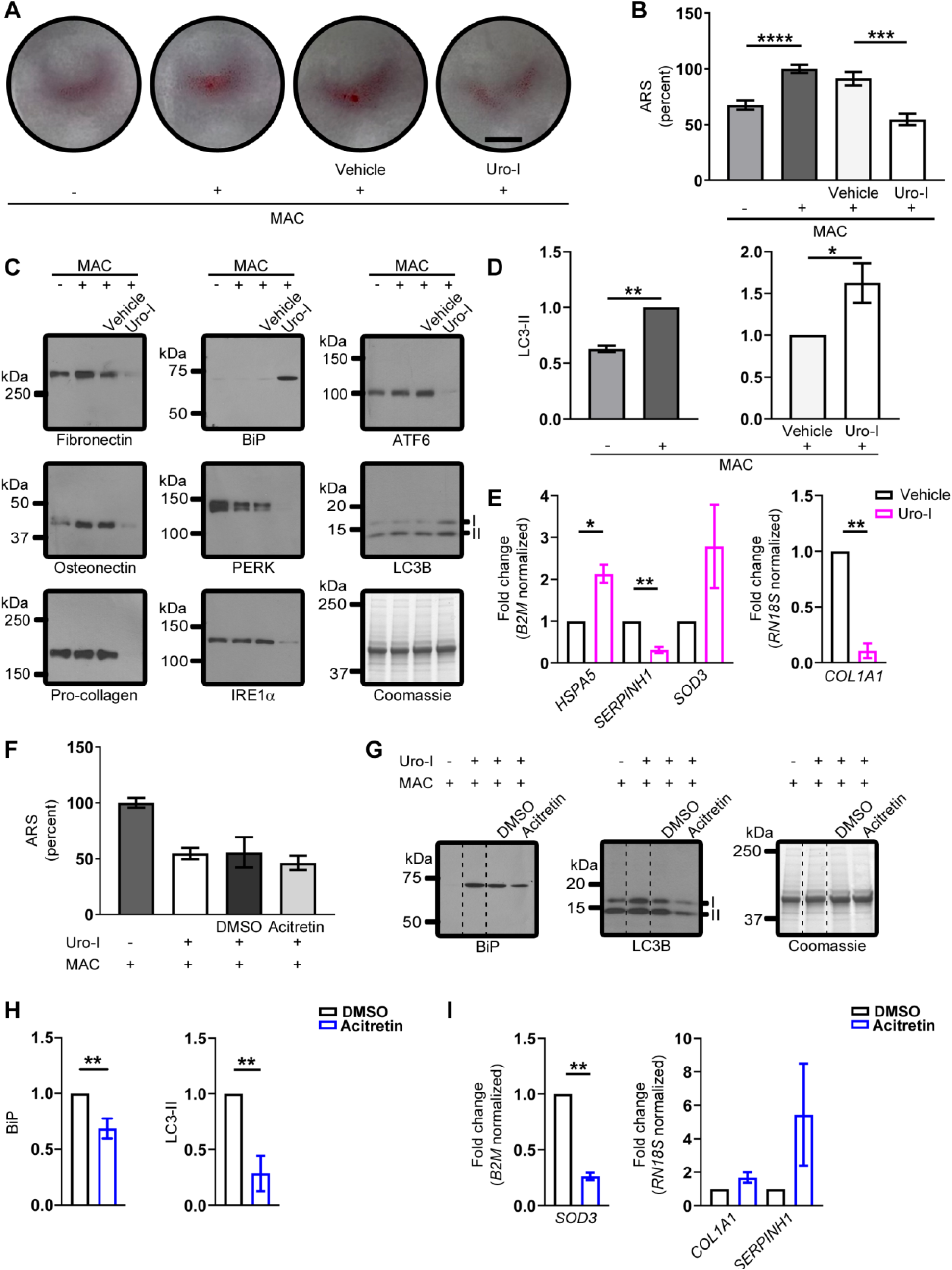
Saos-2 cells mimic CEP zebrafish model. **(A,B)** Mineralization in Saos-2 cells treated with MAC±Uro-I was assayed using ARS staining (photograph, A; quantification, B). Staining was normalized to MAC only-treated cells (set to 100%). **(C)** Cell lysates from experiment in (A) were blotted with the indicated antibodies. **(D)** Quantification of LC3-II shown in (C). LC3-II level was normalized to MAC only (left panel) or vehicle-treated (right panel), set to 100%. **(E)** RT^2^ Profiler PCR Array (left panel) and qPCR (right panel). Relative gene expression is represented as fold change normalized to housekeeping gene. Data are from 2 independent experiments. **(F)**Acitretin does not rescue reduced mineral matrix phenotype in uro-I-treated cells. ARS staining quantification as in (B). **(G)** Acitretin normalizes ER stress (BiP) and autophagy (LC3-II) markers. Dashed lines represent non-adjacent lanes in the gel. Coomassie-stained gel (C,G) shows equal protein loading. **(H)** Quantification of LC3-II. LC3-II level was normalized to DMSO-treated cells set to 100%. **(I)** Gene expression profiling as in (E). \**p*<0.05, \*\**p*<0.01, \*\*\**p*<0.001, \*\*\*\**p*<0.0001

Since fluorescent porphyrins cause protein aggregation or loss of antibody reactivity when tested by immunoblotting, we tested whether uro-I-mediated inhibition of Saos-2 mineralization led to aggregation of bone matrix proteins. Blotting Saos-2 cell lysates prepared from uro-I or vehicle treated cells using antibodies to fibronectin, osteonectin and type 1 pro-collagen showed a distinct loss of monomer for these proteins after uro-I treatment (Fig.3C). We attribute the loss of antibody reactivity to epitope masking after uro-I binding and subsequent oxidation and aggregation, as shown previously for PP-IX^13^. The loss of matrix protein monomers and aggregation was verified by mass spectrometry (Fig.S3).

Given the effect of uro-I on protein aggregation, we tested whether uro-I treatment initiates unfolded protein response (UPR) and endoplasmic reticulum (ER) stress. We observed upregulation of BiP, consistent with UPR and ER stress^21^ (Fig.3C). Other ER stress markers, including PERK, IRE1α and ATF6, were likely oxidized and aggregated, as judged by monomer loss (Fig.3C). Our findings suggest a non-canonical form of ER stress, which we have observed upon PP-IX accumulation^13^, that involves aggregation and possibly inactivation of ER resident proteins and chaperones.

Autophagy modulates exocytosis of hydroxyapatite crystals and thus plays an important role in bone mineralization by osteoblasts^22,23^. Since uro-I inhibited mineralization, we tested whether it also disrupted autophagy. As expected, MAC-treated Saos-2 cells showed increased LC3-II (Fig.3D, left panel). Uro-I treatment also increased LC3-II levels (Fig.3D, right panel). The likely explanation for the increased LC3-II is not increased autophagy but a slowing of autophagic flux, which could lead to stalling of exocytosis of mineral-loaded vesicles and decreased mineralization of bone matrix^24^.

To further characterize uro-I-mediated impairment of mineralization in Saos-2 cells, we performed gene expression analysis to probe for alterations in the stress response pathway. Genes that were differentially regulated two-fold or more after uro-I treatment were assessed further. Uro-I treatment increased *HSPA5* (2.1x) and *SOD3* (2.8x), while *SERPINH1* decreased (3.3x) (Fig.3E, left panel). Since *HSPA5* encodes BiP, *HSPA5* upregulation supports the BiP upregulation observed biochemically (Fig.3C). *SERPINH1*, a collagen-specific chaperone, downregulation may account for collagen misfolding and aggregation. Because type 1 collagen is the most abundant protein in bone matrix^25^, we assessed *COL1A1* expression and observed a 90% reduction in *COL1A1* after uro-I treatment (Fig.3E, right panel). This finding supports uro-I-induced loss of mineralization, since collagen serves as a matrix for mineral deposition.

We next asked whether acitretin can protect from the effects of uro-I, by treating Saos-2 cells with uro-I in the presence of acitretin. Although acitretin did not prevent uro-I-mediated loss of mineralization (Fig.3F), it blunted the ER stress response by reducing BiP level and normalized the autophagic flux by reducing LC3-II (Fig.3G,H). Acitretin also downregulated *SOD3* 3.8-fold, thereby suggesting that acitretin mitigates the oxidative stress caused by uro-I (Fig.3I). Upregulation of *COL1A1* (1.7x) and *SERPINH1* (5.4x) was also observed.

Taken together, our data demonstrate that acitretin mitigates uro-I-mediated proteotoxicity and oxidative stress. However, under the conditions tested, acitretin did not rescue the impairment of mineralization caused by uro-I treatment of Saos-2 cells. A possible explanation for why mineralization was not normalized by acitretin is that ER stress and autophagy pathways need to be normalized in order for cells to have their mineralization ability restored. Alternatively, acitretin may act differently on various cell types which is one major advantage offered by the *in vivo* zebrafish system.

## DISCUSSION

Uro-I is a fluorescent porphyrin capable of types I/II-photosensitized reactions^26–28^, which explains the observed photosensitivity in CEP, damage to digits and facial features^3^. However, light is unlikely to reach deep internal tissues, which are also affected in CEP. Of note, we observed uro-I-mediated protein aggregation and decreased mineralization in the dark. Previous studies had also reported dark effects of porphyrins. For example, uro-I increased collagen biosynthesis in human skin fibroblasts^29^, and inhibited erythrocytic uroporphyrinogen decarboxylase activity^30^. A 2-hit model could explain light-independent porphyrin-mediated protein aggregation and proteotoxicity whereby, in absence of light, a secondary oxidant source (eg. inflammatory cells) causes protein oxidation followed by porphyrin binding to oxidized protein, yielding protein aggregates^11,12^ (Fig.4). CEP is frequently associated with superinfections and osteolysis^3,31^. Hence, infiltrating immune cell-generated oxidants might serve as a secondary source of oxidant, leading to uro-I mediated protein aggregation in internal organs such as bones. Additionally, uro-I might generate oxidants by acting as a substrate for the ferredoxin/ferredoxin:NADP+ oxidoreductase system^32^. Although ferredoxin/ferredoxin:NADP+ oxidoreductase are commonly associated with hepatic microsomes, they are also expressed in osteoblasts^33^ and could metabolize uro-I to generate oxidants in the absence of light.

**Figure 4.**
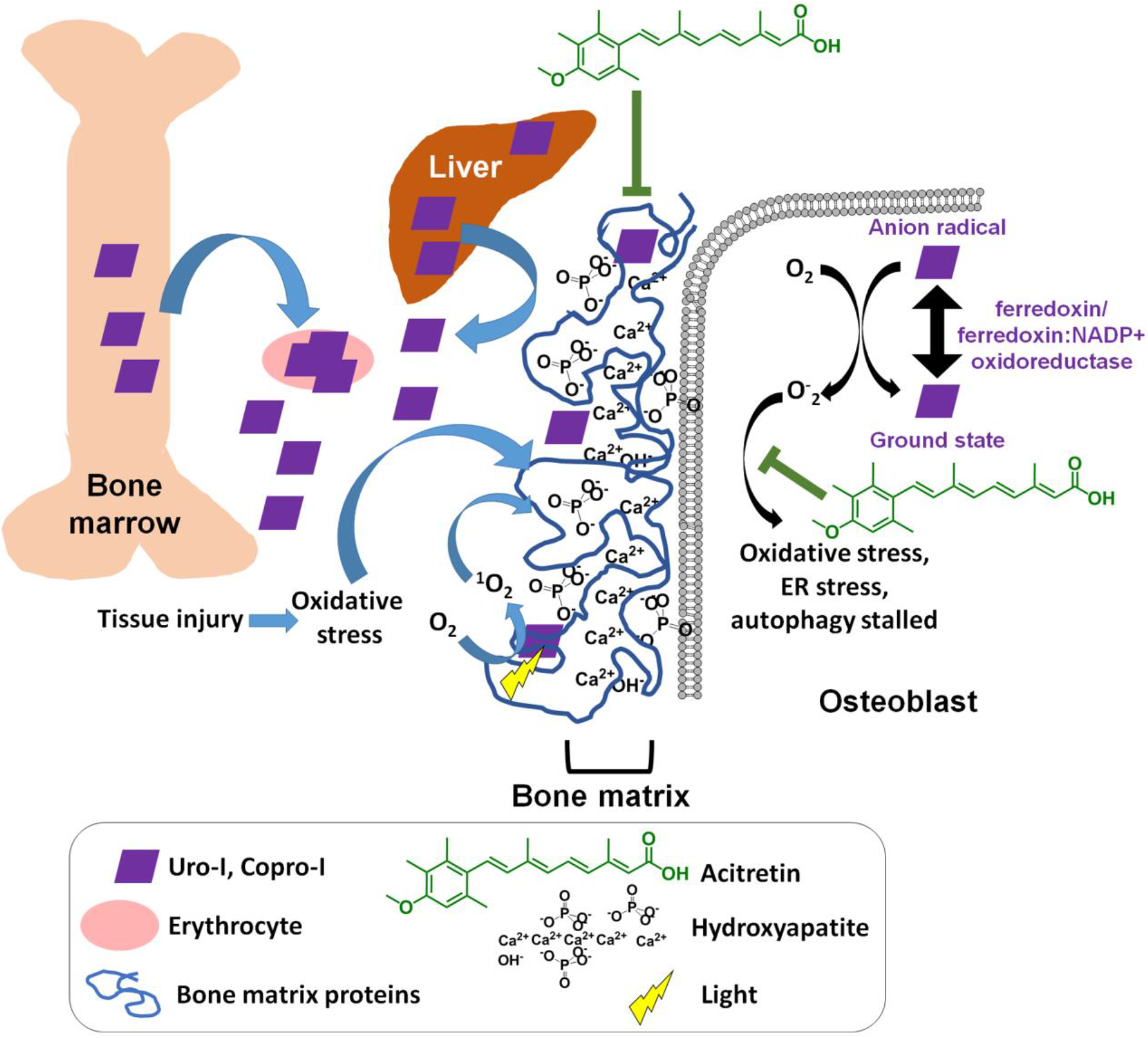
Proposed model of CEP pathogenesis. UROS inhibition leads to production of uro/copro-I mostly in erythrocytes and liver, which is transported through blood to the bones. Uro-I causes bone damage by binding to hydroxyapatite, causing oxidative and ER stress, protein aggregation and stalled autophagy. Acitretin partially rescues uro-I-induced bone damage by reducing oxidative and ER stress and restoring autophagic flux.

The differences in charge and polarity of uro-I and PP-IX might explain the striking difference in their tissue localization. Retro-orbitally injected PP-IX accumulated in zebrafish liver^10^, while uro-I accumulated preferentially in bone (Fig.1). Of note, liver cancer cell lines do not uptake uroporphyrin (unpublished data), possibly due to its high negative charge that prevents traversing the cell membrane^34^. Based on our data, we propose that negatively charged uro-I binds to Ca^2+^ in hydroxyapatite (Fig.4) and thus bone and Saos-2 cells are affected by uro-I. This association with bone matrix causes uro-I to have a different protein aggregation signature compared to PP-IX, which is primarily internalized. PP-IX aggregated intracellular proteins such as keratins and glyceraldehyde 3-phosphate dehydrogenase^13,15,35^, whereas uro-I affected extracellular bone matrix proteins (Fig.3). Oxidants such as singlet oxygen, a major oxidant produced by photosensitive reactions^26^, have extremely small intracellular diffusion distance (10-20nm) and lifetime (10–40ns)^36–38^. Binding of uro-I to bone matrix causes a ‘sensitizer-acceptor’ coupling, as observed for other diffusible oxidants^39,40^, and greatly increases the oxidation efficiency and specificity. Of note, oxidized fibronectin reduces mineralization of rat calvarial osteoblasts *in vitro*^41^. The high selectivity of uro-I localization to bone matrix might provide a pathway to develop photodynamic therapeutic agents for bone cancers such as osteosarcoma.

The management of CEP is challenging, with current therapeutic options focusing on bone marrow/hematopoietic stem cell transplantation^3^, and by avoidance of sun and light exposure, including the use of protective clothing^42–45^. There are also potential experimental therapeutic approaches including gene therapy^46^, proteasomal inhibitors^47,48^, iron chelation^49,50^ and phlebotomy^51^. Most recently, the repurposed use of ciclopirox, an approved broad-spectrum antifungal agent, showed promising results in the treatment of CEP using a mouse model^52^. However, there are limitations to these approaches, such as complications from transplantation and neurotoxic side effects of proteasome inhibitors^53^. Currently there are no known pharmaceuticals that act by clearance of uro-I, and in this regard acitretin provides a novel approach. Acitretin might also act as an antioxidant (Fig.4) due to its hyperconjugated nucleophilic double bonds. Thus, through a combination of destabilizing uro-I-bone matrix interaction and antioxidant activity, acitretin could ameliorate CEP manifestations (Fig.4). Acitretin also offers a drug repurposing advantage since it is already approved for psoriasis treatment^54^.

## MATERIALS AND METHODS

### Zebrafish experiments and cell culture

Zebrafish (*Danio rerio*) experiments were conducted using ABxTL hybrid and NHGRI-1 wild type zebrafish lines. All animal procedures were approved by the Rutgers University Institutional Animal Care and Use Committee (protocol number PROTO201900147) and performed in compliance with federal guidelines and the standards of the NIH Guide for the Care and Use of Laboratory Animals^55^, the Rutgers University IACUC Policy Handbook and the Animal Research: Reporting of *In Vivo* Experiments (ARRIVE) guidelines.

Saos-2 cells were purchased from ATCC. Cells were maintained in McCoy 5A medium supplemented with 15%FBS, penicillin/streptomycin, non-essential amino acids, Hepes and L-glutamine. To induce mineralization, cells were treated with mineralization activation cocktail (MAC), consisting of 5mM β-glycerophosphate, 50μM ascorbic acid and 10nM dexamethasone^56^.

### Uro-I solution preparation and treatment of zebrafish larvae and Saos-2 cells

Uro-I (uroporphyrin-I dihydrochloride; Frontier Scientific, Catalog#:U830-1) was initially resuspended in 0.1M NaOH and the pH was adjusted to neutral using 0.2 M Na_2_HPO_4_. Six days post fertilization (dpf), ABTL zebrafish larvae were injected via the retro-orbital route with approximately 3nL of 7.2mM Uro-I solution and control larvae were injected with vehicle (0.1M NaOH in 0.2M Na_2_HPO_4_). After injection, larvae were immediately transferred to Petri dishes wrapped with heavy duty aluminum foil and kept in a dark incubator, at 28.5°C for 24h. Where indicated, 7 dpf larvae were injected with approximately 2 nL of 0.2% w/v calcein (Sigma, Catalog#:C0875) 2h prior to imaging.

Saos-2 cells were plated in 12-well plates (1.5×10^5^ cells/well) and allowed to attach overnight. Cells were then treated with Uro-I (144μM final concentration) or vehicle in medium containing MAC for 3 days. Experiments were conducted in a dark room and cells were kept shielded from light in a tissue culture incubator.

### Confocal microscopy imaging and quantification

Seven dpf ABxTL zebrafish larvae were anesthetized with tricaine-S (Syndel) and immobilized in 0.5% low melt agarose. Fluorescent z-series were captured using an Olympus FV500 confocal microscope (10X objective, confocal aperture of 300μm) with an optical thickness of 10μm and z-step size of 10μm. Calcein was excited with a 488nm argon laser and emission was captured between 505 and 525nm. Porphyrin was excited with a 405nm laser diode and emission was captured above 560nm. Three-dimensional image reconstruction and quantification of fluorescent signal and bone volume were performed using Imaris 3D visualization and analysis software v7.7 (Bitplane).

### *In vivo* and *in vitro* binding of uro-I

Six dpf ABxTL zebrafish larvae were injected with Uro-I or vehicle (as described above) and 24 h later were euthanized by tricaine-S overdose on ice bath. Bones were harvested as previously described^57^. Briefly, soft tissue was removed by incubating larvae with Accumax solution (MilliporeSigma, Catalog#:A7089) under vigorous shaking. Bones were collected using a 70μm cell strainer, followed by demineralization with 1.2M HCl. Fluorescent images were captured prior to and after the demineralization step using a Zeiss Axio Imager M2 fluorescence microscope. Porphyrin signal was captured using the red fluorescent channel.

10mg hydroxyapatite (Acros Organics, Catalog#:1306-06-5) was incubated with 1mM Uro-I, 1 mM Copro-I (coproporphyrin-I dihydrochloride, Frontier Scientific, Catalog#:C654-1), vehicle or 0.2% calcein for 30min in the dark and vortexed every five minutes. Samples were washed and imaged by epifluorescence microscopy as described above.

### High throughput drug screening

Unbiased high throughput drug screening was performed using the Prestwick library (Prestwick Chemical), which consists of 1,280 small molecules chosen by the manufacturer for their bioavailability and safety. A pooled approach, where four compounds were tested together, was used in order to optimize animal use and investigation of drugs with potential for CEP treatment. Zebrafish E3 medium (100 μL/well) was transferred to a 96-well half area imaging plate (Corning, cat. n. 3880) using a Multidrop dispenser (ThermoFisher Scientific). Compounds (0.4μL of 2mM stock) were added to the wells using a multichannel plate handling robot (Biomek FX, Beckman Coulter Life Sciences). This step was performed four times in order to pool four compounds into one well: one 384-well stock plate yielded one 96-well test plate. Control wells contained 1.6μL of DMSO.

Six dpf NHGRI-1 zebrafish larvae were injected retro-orbitally with approximately 2nL of a solution of Uro-I (10mM) and calcein (0.2% w/v). Immediately after injection, larvae were transferred to a 96-well test plate (two larvae in 50 μL of E3 medium/well), including the DMSO control wells. Control larvae injected with the drug vehicle (dimethyl sulfoxide, DMSO) and calcein were transferred to E3 medium-only containing wells. Larvae were kept in the dark, at 28.5°C. After 24h, they were anesthetized with tricaine-S, centrifuged at 500xg for two minutes and imaged using the ImageXpress Micro Cellular Imaging and Analysis System (Molecular Devices). Positive hits were selected based on visual identification of calcein signal increase and porphyrin signal decrease compared to DMSO-treated larvae. Compounds in test wells that met the inclusion criterion were tested individually in the same manner as described above (Figure S2).

### Acitretin validation and treatment

A dose-response curve with acitretin (Selleck Chemicals, Houston, TX) was conducted (0.5-12.5μM) and 10μM was observed to yield consistent results, without being toxic to zebrafish larvae. Validation and characterization of acitretin as a potential treatment for CEP was performed. Six dpf ABTL zebrafish larvae injected with uro-I were immediately transferred to 10cm plastic dishes containing 10μM acitretin or DMSO in E3 medium (prophylaxis protocol, Fig.2A), and incubated for 24 h in the dark at 28.5°C. Porphyrin binding to bones and bone volume were analyzed by confocal microscopy as described above. Porphyrin excretion into the medium was quantified. Uro-I-injected larvae were transferred to 96-well plates, one larva/well, 100μL of 10μM acitretin or DMSO/well. Medium was collected after 24h and porphyrin was quantified as described previously^13^. Etretinate (Selleck Chemicals, Houston, TX) and tretinoin (Selleck Chemicals, Houston, TX) treatment was performed as described for acitretin.

In addition to being used as prophylaxis, we evaluated whether acitretin had a therapeutic effect. Six dpf ABxTL zebrafish larvae were injected with Uro-I and 24h later they were transferred to E3 medium containing 10μM acitretin or DMSO. Porphyrin binding, excretion and bone volume were analyzed as described above. For Saos-2 cells, they were treated with 10μM acitretin or DMSO in medium containing MAC and Uro-I.

### Alizarin Red S (ARS) staining and quantification

Cell mineralization was quantified by ARS (Sigma Aldrich St. Louis, MO) staining as described previously^58^ with minor modifications. Briefly, cells were fixed with 100% ethanol at 37°C for 1h, stained with 40mM (pH4.2) ARS solution for 20min in an orbital shaker. Cells were washed and ARS was extracted by incubation of fixed cells with 10% (v/v) acetic acid, followed by scraping, incubation of suspension (85°C, 10min), centrifugation and neutralization of supernatant with 10% (v/v) ammonium hydroxide. ARS standard curve (from 2-0.02mM) and samples were transferred in triplicate to a 96-well plate and absorbance was measured at 405nm.

### Cell harvest, immunoblotting and mass spectrometry

Saos-2 cells were lysed in ice cold RIPA buffer (Sigma Aldrich, St. Louis, MO) with protease inhibitor cocktail (Thermo Scientific, Waltham, MA) and scraped. Whole cell lysate was kept in the dark until reducing SDS-PAGE sample buffer was added. Immunoblotting, band densitometry and mass spectrometry were conducted as described previously^12,13^. The antibodies used and their vendors are as listed. Antibodies to the indicated antigens (and sources) are: ATF6, BiP, LC3B (Cell Signaling Technology, Danvers, MA); fibronectin HFN 7.1, pro-collagen SP1.D8, osteonectin AON-1 (Developmental Studies Hybridoma Bank; Iowa City, Iowa); IRE1α, PERK (Invitrogen, Carlsbad, CA); lamin A/C (Santa Cruz Biotechnology, Dallas, TX).

### Gene expression profiling

Saos-2 cells RNA was extracted using RNeasy mini kit (Qiagen, Catalog#:74104). and gene expression was carried using the RT² Profiler PCR Array for human cellular stress responses (Qiagen, Catalog#:PAHS-019ZA) following manufacturer’s instructions. A previously described qPCR^59^ was performed for *COL1A1* and *SERPINH1* (IDT Integrated DNA Technologies, PrimeTime assay ID Hs.PT.58.15517795 and Hs.PT.56a.26865778, respectively).

### Statistical analysis

Statistical analysis was performed using GraphPad Prism v8 (GraphPad Software). Unpaired two-tailed Student’s t-test was used to determine statistical significance. Error bars represent standard error of the mean. \**p*<0.05, \*\**p*<0.01, \*\*\**p*<0.001, \*\*\*\**p*<0.0001.

## ACKNOWLEDGEMENTS

We thank the following research cores at the University of Michigan: Center for Chemical Genomics; Microscopy, Imaging and Cellular Physiology Core (MICPC) Imaging Laboratory; and the Proteomics Research Facility.

## AUTHOR CONTRIBUTIONS

Conceptualization: J.B.C., J.S.E., D.M., M.B.O.; Methodology: J.B.C., J.S.E., J.A.S.; Investigation: J.B.C., J.S.E., N.K., R.A.D., A.C.F., M.S.G., S.I.L.; Writing original draft: J.B.C., D.M.; Review and editing of the manuscript: J.B.C., D.M., M.B.O.; Review of final version prior to submission: all authors; Overall Project Supervision: M.B.O.; Funding acquisition: M.B.O., J.A.S.

## SUPPLEMENTARY FIGURES

**Figure S1.**
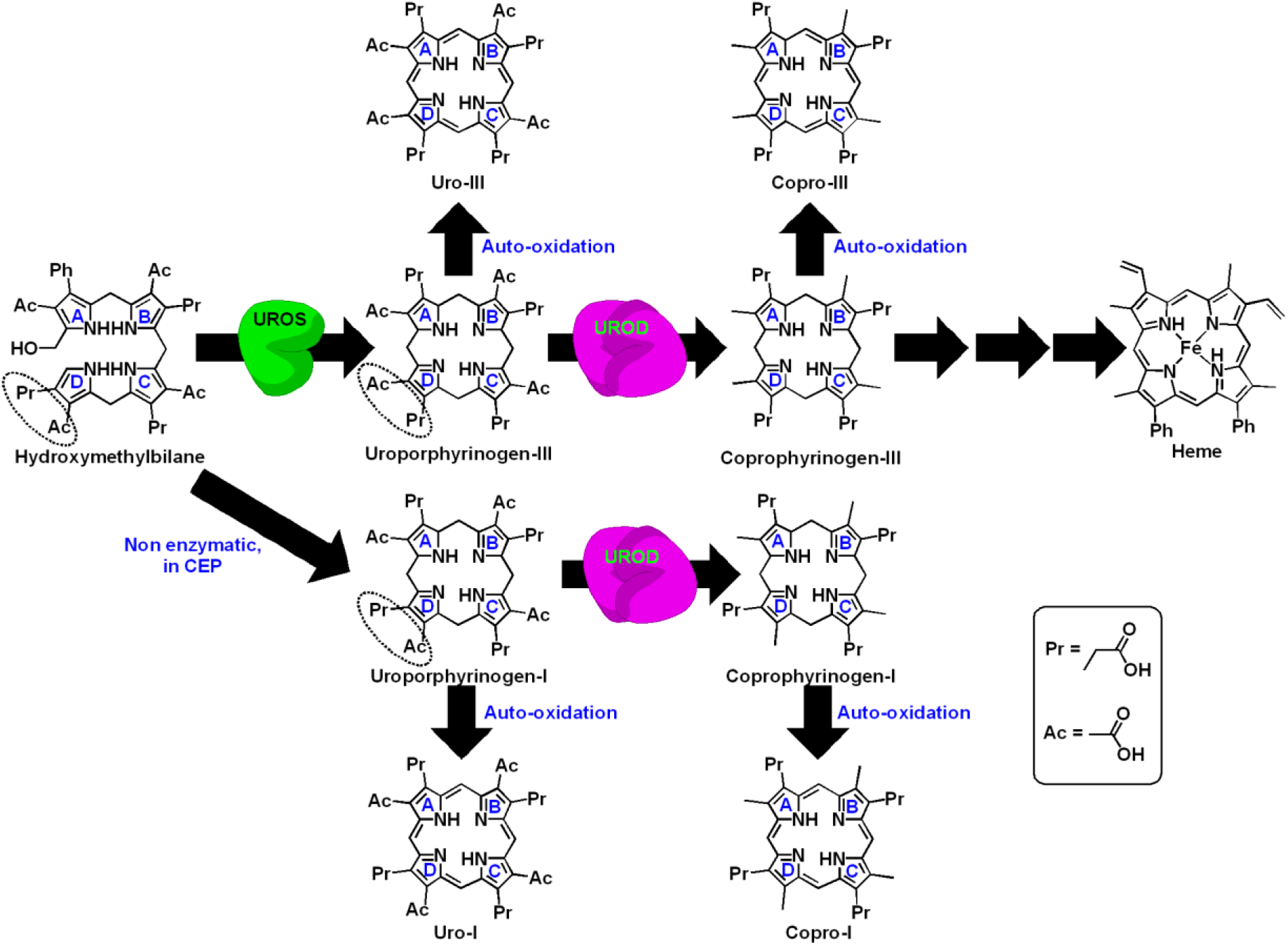
Uroporphyrinogen III synthase (UROS) inhibition accumulates uro-I and copro-I in CEP. UROS, a cytosolic enzyme, catalyzes the conversion of the linear tetrapyrrole, hydromethylbilane (HMB) to the first cyclic tetrapyrrole of the pathway, uroporphyrinogen-III^1,2^. UROS ‘flips’ the position of the acetate and propionate in the ‘D’ pyrrole ring and subsequently causes ring closure to form uroporphyrinogen-III (dotted oval)^1,3^. Uroporphyrinogen-III is decarboxylated by uroporphyrinogen decarboxylase (UROD) to form coproporphyrinogen-III, which through a multi-step mechanism that involves the formation of protoporphyrin-IX, generates heme. In absence of UROS activity, there is spontaneous ring closure of HMB to form uroporphyrinogen-I, a positional isomer of uroporphyrinogen-III, where the acetate/propionate inversion in ring ‘D’ does not occur. Uroporphyrinogen-I is decarboxylated by UROD to coproporphyrinogen-I, but after this step the pathway gets blocked since coproporphyrinogen-I cannot be metabolized by coproporphyrinogen oxidase. Porphyrinogens are relatively unstable compounds, and are auto-oxidized from their colorless, non-fluorescent porphyrinogen forms to colored, fluorescent porphyrins^4^. Thus UROS blockade leads to accumulation of uroporphyrin-I (uro-I) and coproporphyrin-I (copro-I).

**Figure S2.**
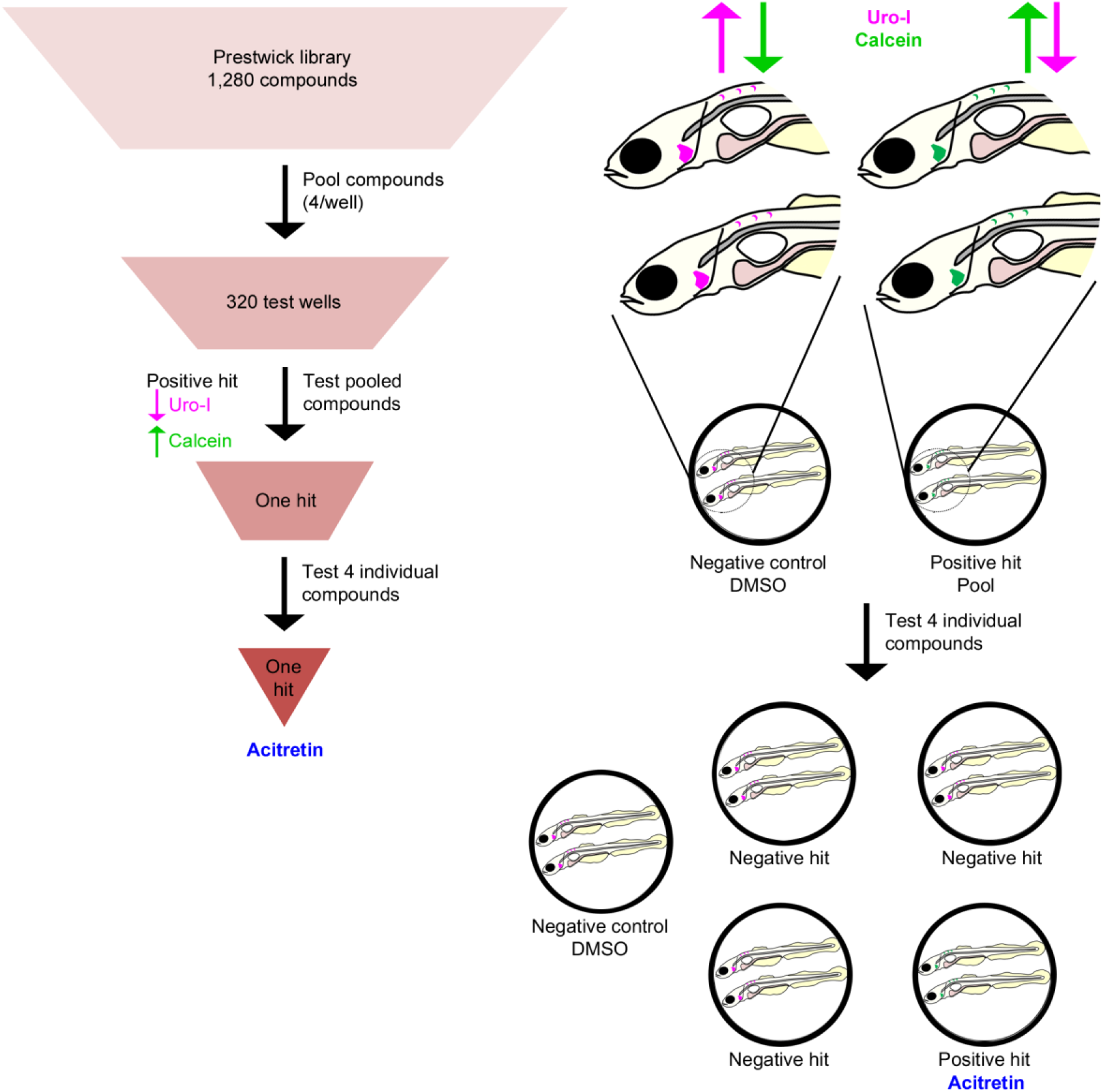
High throughput drug screening for CEP. High throughput drug screening protocol to identify potential drug treatments for CEP was conducted by testing 1,280 small molecules from the commercially available Prestwick library. Initial screening was performed by pooling four drugs per well, with two zebrafish larvae in each well. 6dpf zebrafish larvae were injected with uro-I and calcein simultaneously. 24h later, they were imaged by epiflourescence microscopy using the automated ImageXpress system. Visual analysis was conducted and identification of wells containing larvae with reduced uro-I and increased calcein signal (magenta and green arrows, respectively) in bones compared to DMSO-treated larvae were selected for individual testing of each drug. Of the 320 pools tested, one was identified as potential hit. Once the four drugs were tested individually, acitretin was identified for decreasing uro-I accumulation in bones.

**Figure S3.**
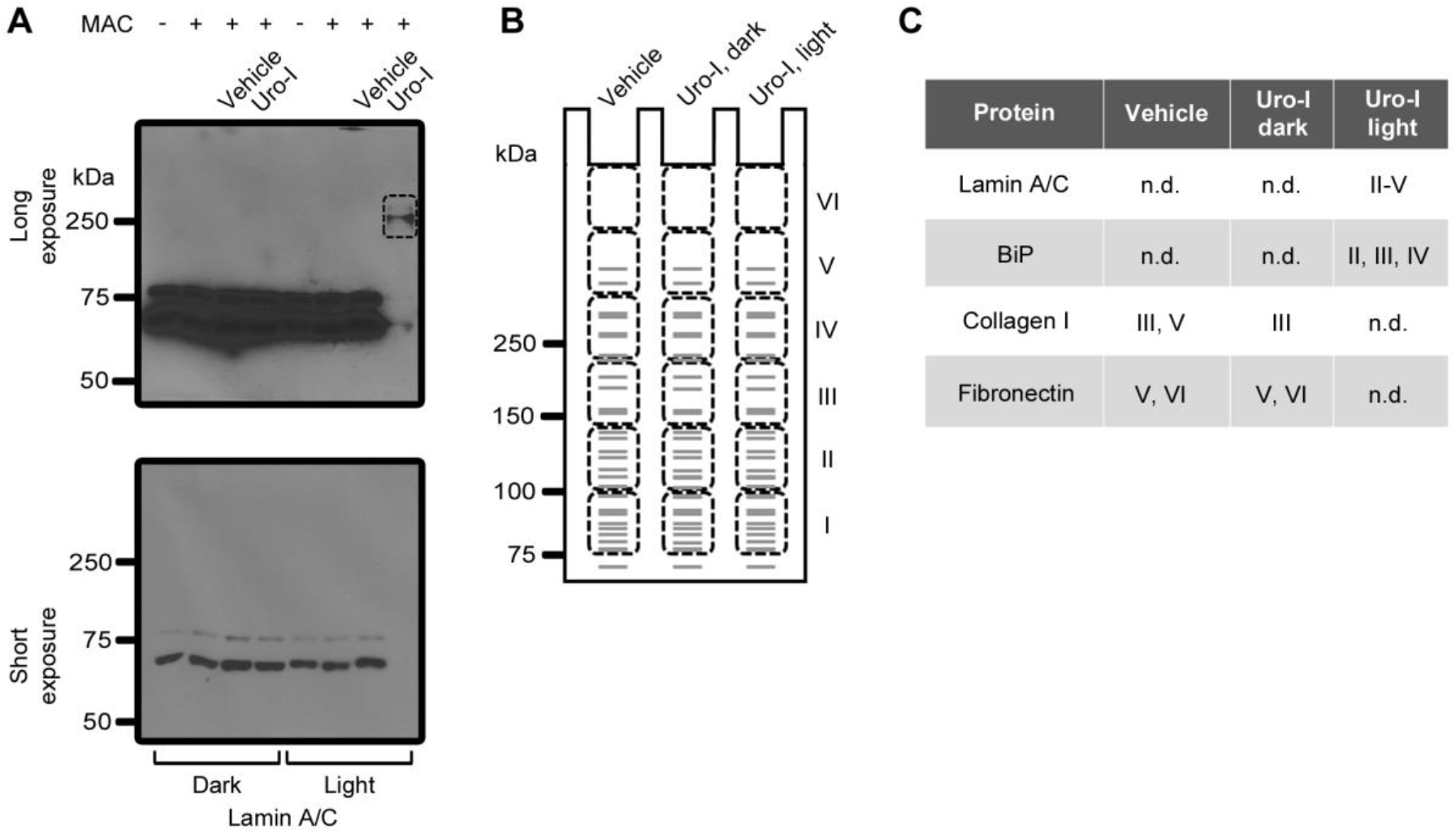
Uro-I causes aggregation of bone matrix proteins in a light-independent manner. **(A)** Saos-2 cells were treated for three days with uro-I or vehicle in the presence of mineralization activation cocktail (MAC). Cells grown in medium without MAC (no mineralization stimuli) and in MAC alone were used as controls for MAC efficiency. Experiments were performed in a dark room and cells were shielded from light throughout the whole experiment. In order to verify whether protein aggregation took place while cells were alive and represented a biologically relevant finding, or if aggregation was an artifact of light exposure during processing of samples, an aliquot of lysate from uro-I treated cells was exposed to light prior to addition of reducing SDS-PAGE sample buffer, which we have shown previously that prevents light-induced protein aggregation by porphyrins in cell lysate. Uro-I treatment did not cause lamin A/C to aggregate, with monomer being comparable between vehicle- and uro-I treated cells (3^rd^ and 4^th^ lanes, short exposure). However, upon light exposure of the uro-I treated cells lysate, loss of monomer and high molecular aggregates were observed (7^th^ and 8^th^ lanes, long exposure). These findings confirm that accidental light exposure of samples did not happen, and any protein aggregation observed was a true biological event, not an artifact of cell processing. **(B)** We conducted a proteomics experiment of cell lysates treated with uro-I and vehicle in the dark to further confirm our findings that bone matrix proteins aggregated upon uro-I treatment. Six 1cm regions of a coomassie stained gel (I-IV, cartoon) spanning from the bottom of the well to slightly above the 75kDa marker were cut and submitted to mass spectrometry analysis. **(C)** Our results confirmed lamin A/C aggregated only in the light-exposed uro-I treated cells lysate, but not in vehicle or uro-I treated cells lysate processed in the dark. Furthermore, data revealed that BiP only aggregated as an artifact of light exposure, not in living cells. Lamin A/C and BiP monomers were not detected in the mass spectral analysis because the gel blocks cut did not include the region where lamin A/C and BiP monomers migrate. Lastly, collagen type I alpha I chain and fibronectin were less abundant in uro-I treated cells lysate processed in the dark compared to control (data not shown). Interestingly, there was no collagen or fibronectin detected in the light processed cell lysate. This confirms that loss of monomer is a reliable read out for protein aggregation and that bone matrix proteins are likely forming high molecular weight aggregates that are unable to migrate into the gel.

